# Dissection of Major Depressive Disorder using polygenic risk scores for Schizophrenia in two independent cohort

**DOI:** 10.1101/054973

**Authors:** HC Whalley, MJ Adams, LS Hall, T-K Clarke, AM Fernandez-Pujals, J Gibson, E Wigmore, Jonathan Hafferty, SP Hagenaars, G Davies, A Campbell, C Hayward, SM Lawrie, DJ Porteous, IJ Deary, AM McIntosh

**Affiliations:** Division of Psychiatry, University of Edinburgh, Edinburgh, UK; Centre for Cognitive Ageing and Cognitive Epidemiology, University of Edinburgh, Edinburgh, UK; Department of Psychology, University of Edinburgh, Edinburgh, UK; Centre for Genetics and Molecular Medicine, IGMM, Western General Hospital, University of Edinburgh, Edinburgh, UK; MRCHuman Genetics Unit, IGMM, University of Edinburgh

## Abstract

Major depressive disorder (MDD) is known for its substantial clinical and suspected causal heterogeneity. It is characterised by low mood, psychomotor slowing, and increased levels of the personality trait neuroticism; factors which are also associated with schizophrenia (SCZ). It is possible that some cases of MDD may have a substantial genetic loading for SCZ. A sign of the presence of SCZ-like MDD sub-groups would be indicated by an interaction between MDD status and polygenic risk of SCZ on cognitive, personality and mood measures. In the current study, we hypothesised that higher SCZ-polygenic risk would define larger MDD case-control differences in cognitive ability, and smaller differences in distress and neuroticism. Polygenic risk scores (PGRS) for SCZ and their association with cognitive variables, neuroticism, mood, and psychological distress were estimated in a large population-based cohort (Generation Scotland: Scottish Family Health Study, GS:SFHS). Individuals were divided into those with, and without, depression (n=2587 & n=16,764 respectively) to test whether there was an interaction between MDD status and schizophrenia risk. Replication was sought in UK Biobank (n=33,525). In both GS:SFHS and UK Biobank we found significant interactions between SCZ-PGRS and MDD status for measures of psychological distress and neuroticism. In both cohorts there was a reduction of case-control differences on a background of higher genetic risk of SCZ. These findings suggest that depression on a background of high genetic risk for SCZ may show attenuated associations with distress and neuroticism. This may represent a causally distinct form of MDD more closely related to SCZ.

## Introduction

Major depressive disorder (MDD) has a lifetime prevalence of approximately 16% and is a heritable and severely disabling psychiatric condition (1). It is highly clinically heterogeneous, relying on diagnostic criteria that are fulfilled through the presence of an arbitrary threshold of symptoms that lack empirical validation. Further, individuals passing this threshold can have a very different combination of symptoms. As a likely consequence of causal heterogeneity, progress in our understanding of depression’s underlying risk factors and biological mechanisms has been limited.

MDD is characterised by sustained negative affect, psychological distress and by differences in personality and behavioural traits, such as increased levels of the personality trait neuroticism (2). People with MDD also manifest, on average, reduced executive function, attention, processing speed and working memory (3), but relatively preserved general cognitive ability (4). These behavioural and personality features are partly heritable, enduring features of illness that are often genetically correlated with liability to MDD (5–8). Notably however, these clinical and personality associations are not specific to MDD and also occur in other psychiatric disorders including schizophrenia (SCZ) (9–11).

Genetic factors account for ~37% of liability to MDD and depressive symptoms (12, 13). Genome-wide association studies (GWAS) have indicated that ~30% of the heritability for MDD is accounted for by an additive polygenic genetic architecture, where risk is conferred by an accumulation of many alleles of small effect (14). There is also a genetic overlap with other major psychiatric disorders, including SCZ (15). For example, two papers have shown genetic correlation between MDD and SCZ using both genome-wide association analysis identifying shared risk loci (15), or an approach estimating genetic correlation from sampled SNPs (16). These findings have been interpreted as showing that there are genetic variants that raise liability to both MDD and SCZ. Recently, however, a second possibility has been proposed — that the genetic correlation between SCZ and MDD is caused not by pleiotropy, but by the misclassification of SCZ cases as MDD. This possibility has recently been tested using a novel method aimed at using genotype data to distinguish pleiotropy from heterogeneity, where early evidence suggested that some cases of SCZ may have been misclassified as MDD, though this did not reach significance in this proof of technique study (17). If confirmed, this would imply that some cases of MDD would be a hidden ‘forme fruste’ of SCZ, or that some individuals with MDD may be on a trajectory towards SCZ, but have yet to manifest the clinical phenotype.

Effects of polygenic risk for SCZ have also been demonstrated on a number of related behavioural and cognitive phenotypes. It has been consistently demonstrated for example that polygenic risk for schizophrenia is associated with decreased cognitive ability in healthy control samples (18–21). However, since most studies have examined these relationships in unaffected controls, it is unclear whether the associations between SCZ risk and behaviour/cognitive ability are ameliorated or magnified by the presence of depression. Clarifying this relationship is important because, if MDD consists of a subtype which is closely related to schizophrenia, then that subtype would be expected to show associations with behaviour and cognitive ability that resembled those of SCZ itself.

In the current paper we sought to stratify depression using a continuous scale of SCZ polygenic risk score, hypothesising that case-control differences would depend on genetic risk for schizophrenia, in other words, that there would be a significant interaction between SCZ-PGRS and MDD status. We made two predictions, (1) based on the premise that deficits in cognition are greater in SCZ than in MDD, we first predicted that the high-SCZ-PGRS MDD case-control differences for cognitive variables would be greater compared to those with lower SCZ-PGRS scores. (2) Since neuroticism and psychological distress differ to a greater extent in MDD than SCZ (22) we also predicted that SCZ-PGRS and MDD status interactions would be found for these variables, but that they would be the reverse direction to cognition (attenuated in those with high SCZ-PGRS). We sought to test these predictions in a population based cohort study, the Generation Scotland cohort: The Scottish Family Health Study sample (GH:SFHS) (23, 24). Replication of any significant findings was sought in UK Biobank.

## Materials and Methods

### Main Study Participants

#### Generation Scotland: the Scottish Family Health Study (GS:SFHS)

GS:SFHS is a family-and population-based cohort study. Individuals were recruited at random through general medical practices across Scotland. Initial data collection for GS:SFHS took place between February 2006 and March 2011. The complete study protocol has been described in detail elsewhere (23–25). Details of the mental health and related assessments are summarised below. Ethical approval was provided by NHS Tayside Research Ethics Committee (reference 05/S1401/89). Written consent for the use of data was obtained from all participants.

The full cohort consists of 23,690 individuals who were over 18 years at the time of recruitment and 21,516 of these attended the research clinic. The present study includes 19,351 individuals on whom genome-wide genotype data were available (demographic details presented in Table 1). Pedigree information was available for all participants, and this has subsequently been validated against estimates of relatedness estimated using genome-wide single nucleotide polymorphism (SNP) data on 19,995 individuals.

**Table 1.**
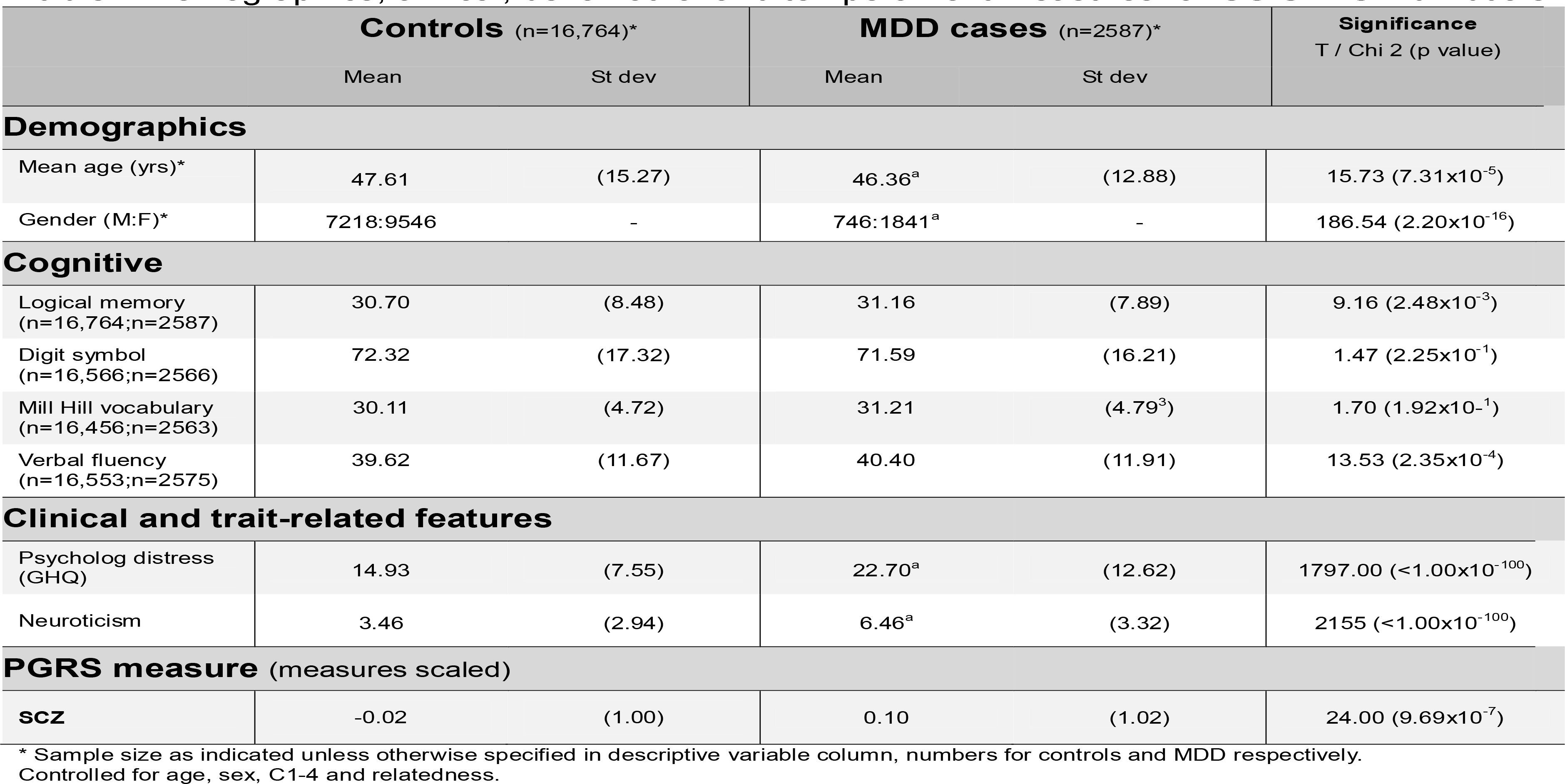
Demographics, clinical, behavioural and temperament measures for GS:SFHS individuals in the current study.

#### Assessment of MDD in GS:SFHS

MDD was diagnosed using the structured clinical interview for the Diagnostic and Statistical Manual of Mental Disorders (SCID) (26). A brief questionnaire was administered to participants to screen for MDD. Participants were asked “Have you ever seen anybody for emotional or psychiatric problems?” and “Was there ever a time when you, or someone else, thought you should see someone because of the way you were feeling or acting?” If they answered yes to either of these questions (21.7% screened positive), they were asked to complete the SCID (26). If they answered no to both of these questions, they were assigned control status. Those who completed the SCID but did not meet the criteria for MDD or other major psychiatric disorder were also defined as controls. Individuals who declined to complete the screening questionnaire or the SCID had their MDD status set to missing. Individuals with a diagnosis of bipolar disorder were removed for the purposes of the current investigation.

#### Cognitive assessment in GS:SFHS

Cognitive abilities were assessed using four tests. Immediate and delayed scores from the recall section of one story of the Wechsler Logical Memory III ^UK^test were summed to provide a measure of verbal declarative memory (27. The Wechsler Digit Symbol Coding test was used to measure processing speed (27). Verbal ability was assessed using the Mill Hill Vocabulary Scale, junior and senior synonyms (28). Executive function was measured using the letter-based phonemic verbal fluency test (letters C, F and L, for one minute each) (29). All test scores were scaled to a mean of zero and standard deviation of one. General cognitive ability (g) scores were obtained by conducting principal component analysis of the tests, and saving the scores on the first un-rotated principal component, on which all tests had substantial loadings (30)

#### Measures of neuroticism and psychological distress GS:SFHS

Measures of psychological distress were assessed using the 28-item General Health Questionnaire (GHQ-28) (31), which consists of 4 subscales designed to assess: (A) somatic symptoms, (B) anxiety and insomnia, (C) social dysfunction and (D) ‘severe depression’. A single measure of global affective symptoms was derived from item responses and the standard Likert scoring totals were used in the current study. The Eysenck Personality Questionnaire-Revised Short Form provided a measure of neuroticism (32). For numbers of individuals completing each measure see Table 2.

**Table 2.**
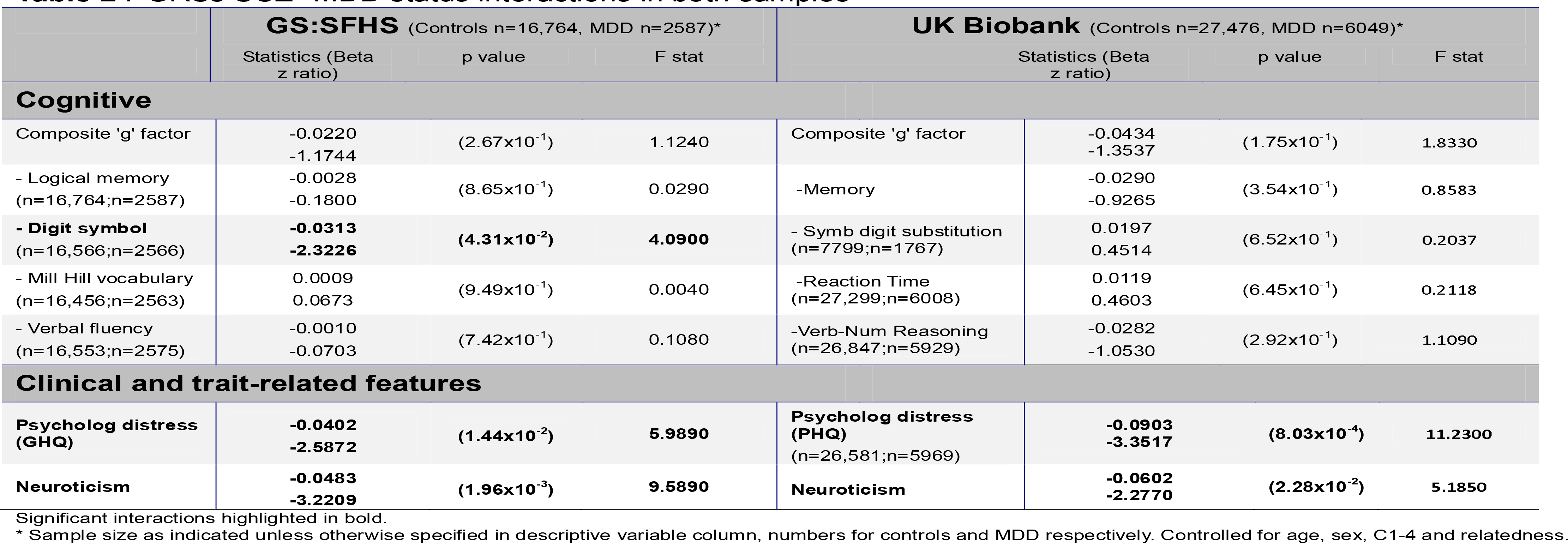
PGRSs SCZ* MDD status interactions in both samples

#### Genotyping and Polygenic Profiling in GS:SFHS

Blood samples were collected using standard operating procedures and stored at the Wellcome Trust Clinical Research Facility Genetics Core, Edinburgh (www.wtcrf.ed.ac.uk) where they were genotyped using the IlluminaHuman0mniExpressExome-8v1.0 BeadChip and Infinium chemistry (33). The genotypes were then processed using the IlluminaGenomeStudio Analysis software v2011.1. The details of blood collection and DNA extraction are provided elsewhere (34).

Polygenic risk scores were calculated using the method described previously (35) using summary data from the most recent GWAS studies from the Psychiatric Genomics Consortium (PGC, 2014). Briefly, SNPs were excluded if they had a minor allele frequency (MAF) < 1%, deviated significantly from Hardy-Weinberg equilibrium (p<1×10^−6^) in the total sample of founder individuals, or had a call rate <99%. Clump based linkage disequilibrium pruning (r^2^ 0.2, 300kb window) was performed to create a SNP-set in linkage equilibrium. Prior to creating polygenic risk scores all strand-ambiguous SNPs were removed from the GS:SFHS dataset and the polygenic risk score for each individual was calculated using the cumulative sum of the number of each SNP allele multiplied by the log of the odds ratio for SCZ across their whole genome. Five SNP set scores were generated using p-value threshold cut-offs of 1, 0.5, 0.1, 0.05 and 0.01. Our primary analyses concerned those SNPs from the PGC data that met a significance level of p=.5 consistent with previous studies (35). Findings for other thresholds are contained within Supplementary Material.

#### Replication sample: UK Biobank

This study includes replication data from the UK Biobank Study (http://www.ukbiobank.ac.uk). UK Biobank received ethical approval from the Research Ethics Committee (REC, reference: 11/NW/0382). UK Biobank is a health research resource that aims to improve the prevention, diagnosis, and treatment of a range of illnesses. Through UK Biobank, 502 655 community-dwelling participants were recruited between 2006 and 2010 in the United Kingdom (36). They underwent cognitive, neuroticism and physical assessments, provided blood, urine, and saliva samples for future analysis, gave detailed information about their backgrounds and lifestyles, and agreed to have their health followed longitudinally. For the present study, genome-wide genotyping data were available on 33,525 individuals. This dataset comprised individuals who survived the quality control process, were independent of individuals within GS:SFHS, who were unrelated, and who did not meet criteria for another major psychiatric disorder. Individuals were assigned to a narrow definition of MDD including cases of moderate or recurrent MDD (see Supplementary Material for further information).

#### Cognitive assessment in UK Biobank

Three cognitive tests were used in the present study to generate measures of general intelligence ‘g’. These tests, which cover the cognitive domains of Reaction Time, Memory and Verbal-numerical Reasoning were collected at the baseline assessment. In addition, data from a second assessment which provided measures of processing speed (similar to the Digit Symbol Coding test, the Symbol Digit Substitution test) were also assessed (further details in Supplementary Material). All test scores were scaled to a mean of zero and standard deviation of one. General cognitive ability (g) was estimated using the principal component analysis approach described above (30).

#### Measures of neuroticism and psychological distress in UK Biobank

In UK Biobank, participants completed the Neuroticism scale of the Eysenck Personality Questionnaire-Revised Short Form (EPQ-R Short Form) (37). As a measure of psychological distress UK Biobank participants undertook the Patient Health Questionnaire (PHQ) which is a multipurpose instrument for screening, diagnosing, monitoring and measuring the severity of depression. As in GS:SFHS scores were scaled to a mean of zero and standard deviation of one.

#### Genotyping and Polygenic Profiling in UK Biobank

Details of the array design, genotyping, quality control and imputation have been described previously (38). Quality control included removal of participants based on missingness, relatedness, gender mismatch, and non-British ancestry. Polygenic profiles were created for SCZ in all genotyped participants using PRSice (39). PRSice calculates the sum of alleles associated with the phenotype of interest across many genetic loci, weighted by their effect sizes estimated from a GWAS of that phenotype in an independent sample. Prior to creating the scores, SNPs with a minor allele frequency < 1% were removed and clumping was used to obtain SNPs in linkage equilibrium with an r^2^< 0.25 within a 200bp window. Multiple scores were then created for each phenotype containing SNPs selected according to the significance of their association with the phenotype. The GWAS summary data was used to create polygenic profiles for SCZ in the UK Biobank participants, at thresholds of p < 0.01, p < 0.05, p < 0.1, p < 0.5 and all SNPs. As in the main GS:SFHS analysis, current results focus on P<0.5.

#### Statistical Analysis

All analyses were conducted in ASReml-R (www.vsni.co.uk/software/asreml, version 3.0). Associations were examined between PGRSs for SCZ and cognition (cognitive factor ‘g’, and contributing tests), psychological distress (GHQ total), and neuroticism (from the Eysenck personality questionnaire) using mixed linear model association analysis in GS:SFHS. Age, age^2^, sex, four multidimensional scaling ancestry components and polygenic profile scores were entered as fixed effects. Since GS:SFHS is a family-based study, to control for relatedness family structure was fitted as a random effect by creating an inverse relationship matrix using pedigree kinship information. SCZ-PGRS and MDD status were entered as main effects in the model and PGRS * MDD status effects were examined by including the interaction term. Where significant interactions were found, further tests were conducted in controls and MDD groups separately. Wald’s conditional F-test was used to calculate the significance of fixed effects. P-values presented are raw p-values uncorrected for multiple testing, results were considered significant if they were reported in both cohorts (p<0.05). Reported beta values are standardised values. The proportion of phenotypic variance explained by polygenic risk score was calculated by multiplying the profile score by its corresponding regression coefficient and estimating its variance. This value was then divided by the variance of the observed phenotype to yield a coefficient of determination between 0 and 1 and converted into a percentage (40).

Significant association and interaction effects were also explored within the independent sample UK Biobank where equivalent measures were available. In UK Biobank polygenic risk profile scores were examined for their association with observed phenotypes and for group interaction effects in ASReml-R using the same methods as above (details in supplementary material), but without the inclusion of a genetic relationship matrix due to the large data set and unrelated nature of the filtered UKB study population used in the current investigation.

## Results

### Demographic, cognitive, trait-related features of MDD and PGRS scores in GS:SFHS

Demographic details are presented in Table 1 for GS:SFHS. Thresholds for case-control group significance were defined as p<2×10^−5^. Groups defined according to depression status were significantly different in terms of sex, psychological distress (GHQ) and for neuroticism (Table 1). The depression group scored higher on the latter two measures. Differences between the groups did not reach significance for the individual cognitive tests. MDD cases and controls also differed significantly on the SCZ PGRS score, with MDD individuals scoring higher.

Demographic details of individuals from UK Biobank included in the current study are presented in Supplementary Table 1. There were significant differences between the groups in terms of age and gender (p<2×10^−5^). The groups also differed according to psychological distress and neuroticism, with the MDD cases scoring higher than controls. Differences between the groups did not reach significance for the individual cognitive tests and MDD cases and controls differed significantly on the SCZ PGRS score, with MDD individuals scoring higher.

### Interaction between MDD status and polygenic risk for schizophrenia, in GS:SFHS and UK Biobank

Results for tests of interaction are presented in Table 2 and Figures 1a,Figures 1b. Significant interaction effects seen across both cohorts were PGRS SCZ * group interactions for measures of psychological distress (β=−0.0402, p=1.44×10^−2^, β=−0.0903, p=8.03×10^−4^, GS:SFHS and UK Biobank) and for neuroticism (β=−0.0483, p=1.96×10^−3^, β=−0.0602, p=2.28×10^−2^, GS:SFHS and UK Biobank). A significant interaction was also reported for measures of processing speed in GS:SFHS (β=−0.0313, p=4.31×10^−2^), but this was not replicated in the UK Biobank sample.

**Figure 1a,b:**
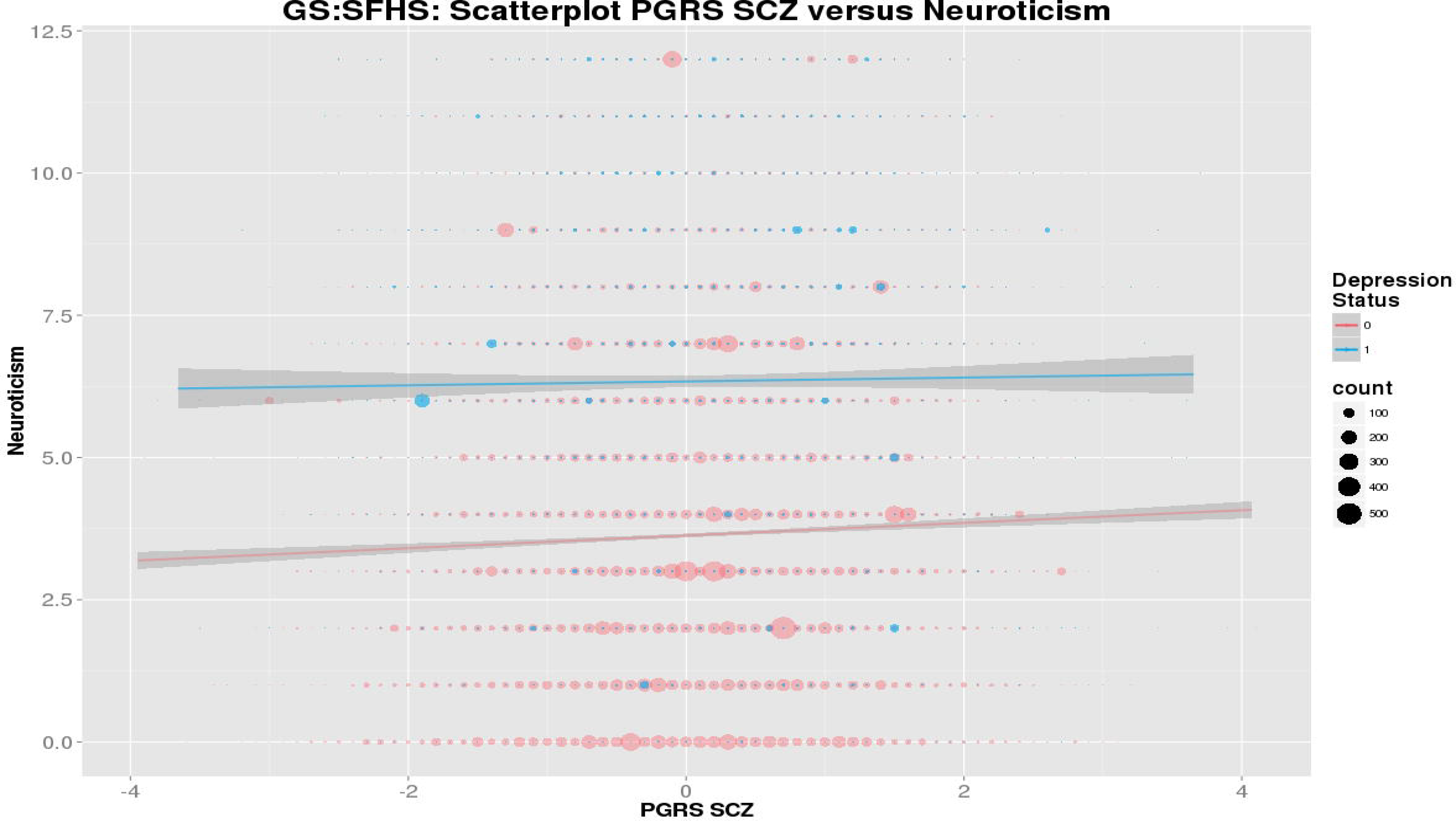
Scatterplot of SCZ-PGRS and neuroticism according to MDD status in GS:SFHS, and UK Biobank respectively.

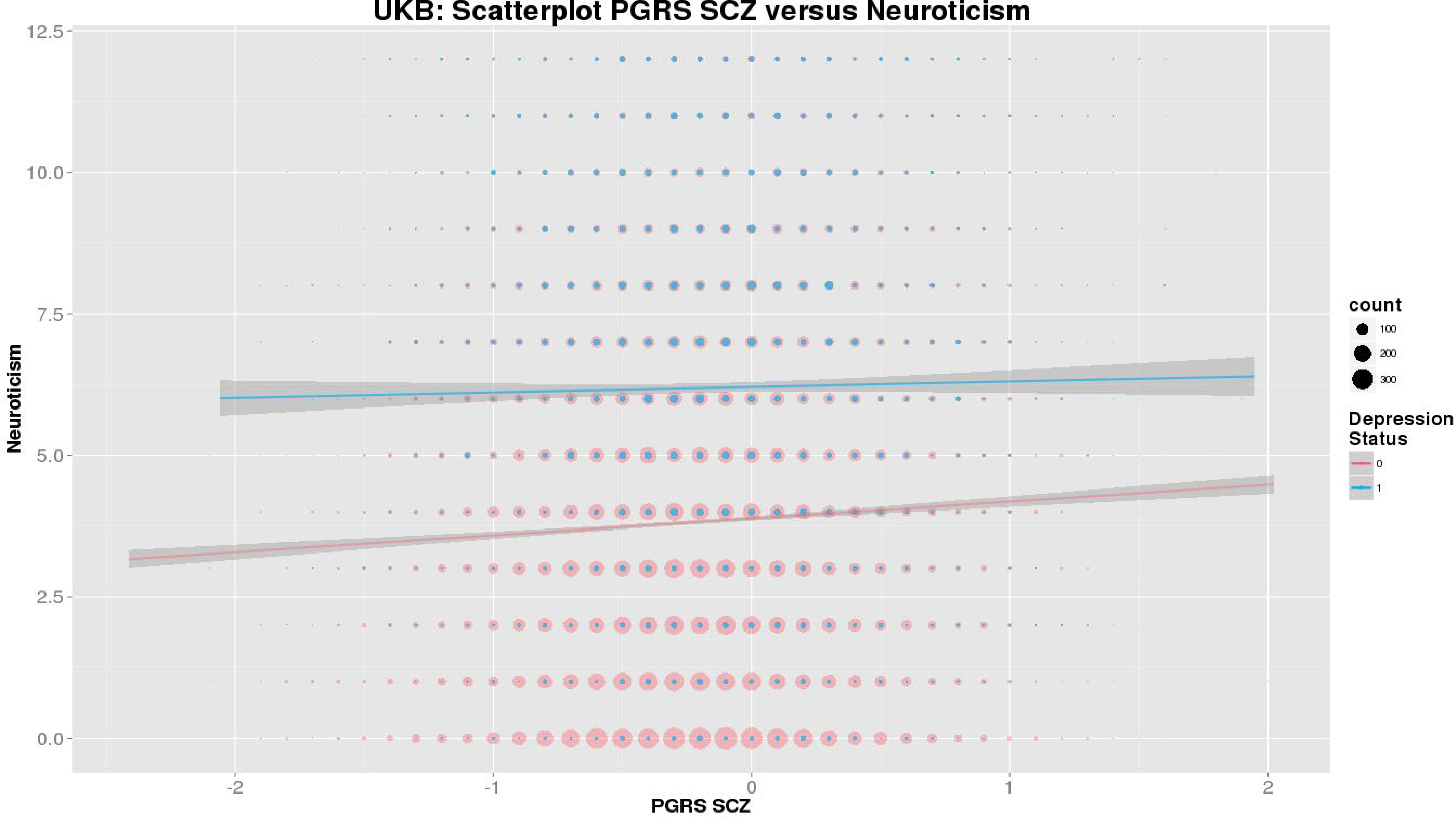

In terms of the direction of associations within diagnostic groups, in both GS:SFHS and UK Biobank samples there was a significant positive association in the control group for measures of psychological distress (β=0.0400, p=1.46×10^−8^, β=0.0827, p=3.25×10^−14^, in GS:SFHS and UK Biobank) and for neuroticism (β=0.0369, p=9.15×10^−6^, β=0.0539, p=4.79×10^−19^, GS:SFHS and UK Biobank) (see Table 3). There were no significant associations within the MDD groups for either measure, in either cohort (see Table 3, Figures 1a,Figures 1b). In terms of psychological distress the proportion of variance explained was 0.15% and 0.22% for control individuals, for GS:SFHS and UK Biobank respectively, and 0.02%, 0.01% for MDD cases. For measures of neuroticism values were 0.17% and 0.30% in controls and <0.01% in MDD cases (see Tables 3 and Tables 4 and Figure 2). For the measure of processing speed in GS:SFHS there was a significant negative relationship in both controls and cases, but the MDD cases demonstrated greater effect size and greater variance explained (1.09%) versus that seen in control individuals (0.33%) (see Table 3). For both distress and neuroticism the SCZ-PGRS * MDD status interactions remained significant after modelling whether individuals were experiencing a current depressive episode at the time of assessment according to the SCID, and when modelling whether individuals were taking antidepressant medication at the time of assessment according to self-report, full details in supplementary material.

**Table 3.**
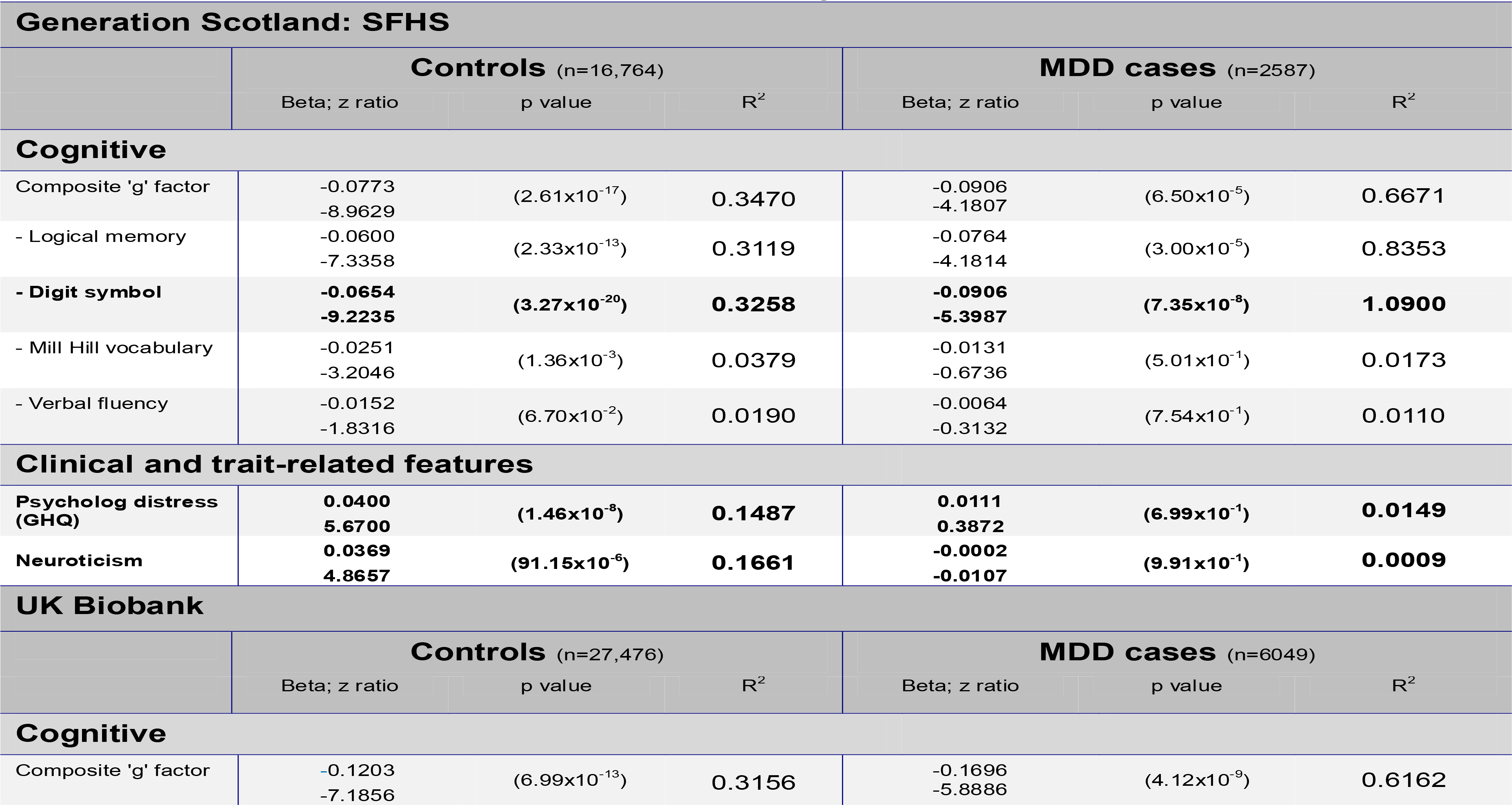
Associations betweenPGRSs for SCZwith cognitive, clinicalandtrait-related features of MDD in cases/controls separately

|  |  |  |  |  |  |  |
| --- | --- | --- | --- | --- | --- | --- |
| -Memory | 0.0657<br>6.3921 | (1.66x10<sup>-10</sup>) | 0.1536 | 0.0490<br>2.2324 | (2.56x10<sup>-2</sup>) | 0.0863 |
| - Symbol digit substitution | -0.1006<br>-5.1415 | (2.79x10<sup>-7</sup>) | 0.2816 | -0.0785<br>-1.9623 | (5.00x10<sup>-2</sup>) | 0.1802 |
| -Reaction Time | 0.0390<br>3.4929 | (4.79x10<sup>-4</sup>) | 0.0422 | 0.0509<br>2.0881 | (3.68x10<sup>-2</sup>) | 0.0692 |
| -Verbal-Numerical Reasoning | -0.1101<br>-9.5051 | (2.16x10<sup>-21</sup>) | 0.3467 | -0.1390<br>-5.5560 | (2.88x10<sup>-8</sup>) | 0.5495 |
| **Clinical and trait-related features** |  |  |  |  |  |  |
| PHQ (health qu) | 0.0827<br>7.5920 | (3.25x10<sup>-14</sup>) | 0.2218 | -0.0259<br>-0.8072 | (4.20x10<sup>-1</sup>) | 0.0111 |
| Neuroticism | 0.0539<br>8.9238 | (4.79x10<sup>-19</sup>) | 0.3026 | 0.0055<br>0.4103 | (6.82x10<sup>-1</sup>) | 0.0029 |

$R^2$ = estimate of variance in trait explained by polygene score in %.

**Figure 2:**
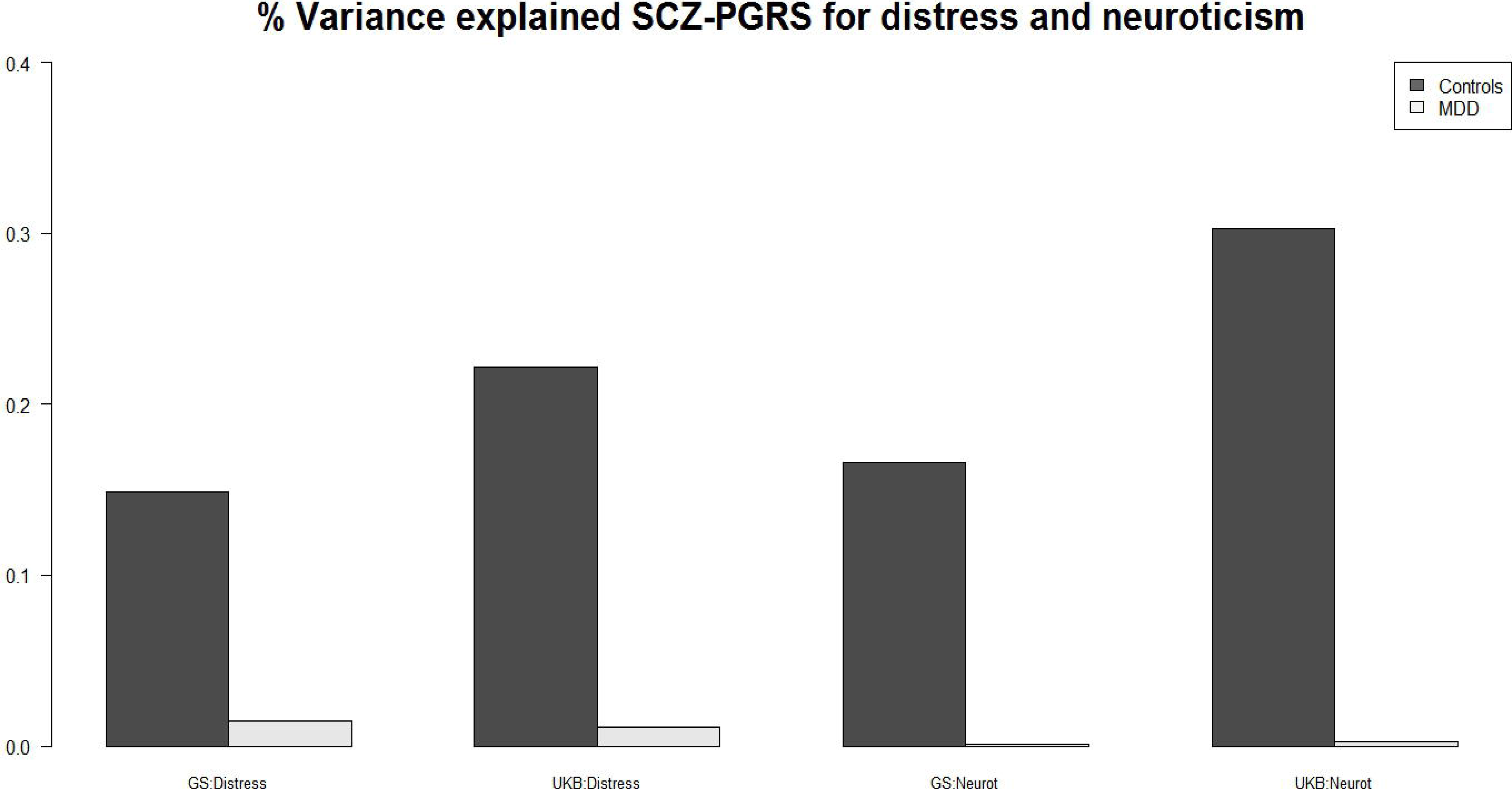
Bar chart of percentage variance explained for distress and neuroticism in GS:SFHS and UK Biobank

## Discussion

Consistent with our predictions we report larger MDD case-control differences in cognition in the context of higher SCZ-PGRS scores in GS:SFHS, although this was not replicated in UKB. We also found MDD case-control differences in neuroticism and distress that were attenuated by higher SCZ-PGRS scores. Lastly, we demonstrate that the greatest variance explained for neuroticism and distress is higher in controls rather than in MDD cases, supporting the suggestion that the MDD group contains a more heterogeneous group comprising differing aetiological subtypes.

### PGRS SCZ associations and interactions with cognition

Previous studies have consistently reported a negative relationship between cognitive function and SCZ-PGRS, indicating shared genetic risk between SCZ and deficits in cognition (18–20, 41). With the exception of one previous study where genetic loading for SCZ (based on proximal family history) was reported to have a substantial negative impact on neurocognition in mood disorder patients (42), we are not aware of any other studies examining this relationship in the context of MDD. In the current study, data from the four individual subtests that contributed to the general factor indicated a significant interaction between SCZ-PGRS and MDD case control status for the digit symbol coding test. This test is a measure of processing speed, which is known to be significantly impaired in SCZ patients, even in relation to other neuropsychological measures (43, 44). Within GS:SFHS, significant negative associations between SCZ-PGRS and processing speed were reported in both controls and MDD cases. In other words, individuals in GS:SFHS with MDD demonstrated greater deficits in tests of processing speed in those who had higher genetic loading for SCZ versus those at low risk. However, the lack of formal replication means these results should be viewed with caution.

### PGRS SCZ associations and interactions with clinical and trait features

Neuroticism is considered to be a stable heritable trait (45) characterized by high tension, irritability, dissatisfaction, shyness, low mood and reduced self-confidence (32). Psychological distress is a more generalised measure of psychiatric well-being or psychological health than a specific psychopathological categorisation. These two traits have been reported to share strong genetic links (7), and there has been considerable support for overlapping genetic risk factors affecting neuroticism and MDD (46). Direct links between genetic risk for schizophrenia and these measures however are less commonly reported than associations with cognitive deficits, as described above.

These measures demonstrated significant interaction effects between SCZ-PGRS and MDD case-control status, seen in both GS:SFHS and UK Biobank cohorts. In addition, the variance explained for these measures was greater in control individuals (~0.20%) than in MDD cases (~0.01%). We suggest that this may be attributable to the aetiologically diverse subgroups within MDD cases. The origins of these interactions were an attenuation of case-control differences on a background of higher genetic risk of SCZ, seen in both cohorts. Neuroticism and psychological distress may therefore be less closely related to the aetiology of depression in the context of high genetic risk for schizophrenia (see Fig 1). It is, however, also possible that the absence of a similar gradient of effect in MDD cases could be due to the substantial effects of depression on distress and neuroticism, leading to a ceiling effect whereby the additional effect of SCZ-PGRS on these phenotypes is undetectable. However, we consider that this is unlikely since the variance of these measures are *not* in fact smaller in the MDD groups in comparison to the control group (which would be the case if there were ceiling effects, see Table 1), and so truncation of these scores at the extreme end seems unlikely. Together, these findings suggest that depression on a background of high genetic risk for schizophrenia may represent a somewhat causally distinct form of the condition.

Although this is a large and arguably well powered study with independent replication, there are important limitations that should be considered. Of note is that the data used to derive polygenic scores do not presently fully account for all heritability attributable to common variation, nor do these measures reflect the contribution of rare variation and copy-number variants. In addition, the degree of variance explained by the SCZ-PGRS score for these phenotypes is relatively modest (0.5-1%). It should be considered however that they are consistent with other previously reported figures of 0-2% for similar phenotypes and in similarly unselected population-based cohorts, and that the higher figures reported in the literature (7-18%) generally relate to the phenotype of SCZ case-control status (see (20)).

Another limitation relates to consistency of testing between these large datasets. Particularly relevant are the differences in administration of the tests of processing speed. Within GS:SFHS, a pen and paper version of the Digit Symbol Coding task from the Wechsler Adult Intelligence Scale III was used, where participants had a two minute time limit to complete the task. In UK Biobank the format of the task was a computerised version where the participant had to click on the number which matched the symbol shown and the participant had one minute to complete the test. Importantly, there were also fewer individuals included in the DSC assessments in UK Biobank (n=7799 controls and n=1767 MDD individuals) than for other cognitive measures, and in relation to GS:SFHS (n=16,566 controls and n=2566 MDD individuals) since this test was conducted at a follow-up assessment rather than at baseline.

In summary, these findings are consistent with a model in which genetic risk for schizophrenia predicts depressive traits in the general population, but that neuroticism and psychological distress may be less closely related to the aetiology of depression on a background of high genetic risk for schizophrenia. This may represent a somewhat causally distinct form of MDD more closely related to SCZ. The study of the genetic basis of variation in such measures is likely to further the understanding of mechanisms by which SCZ genes affect neural function in the context of health and depressive illness.

## Acknowledgements

This study is supported by a Wellcome Trust Strategic Award “Stratifying Resilience and Depression Longitudinally” (STRADL) (Reference 104036/Z/14/Z). Generation Scotland (GS:SFHS) received core support from the Chief Scientist Office (CSO) of the Scottish Government Health Directorates [grant number CZD/16/6] and the Scottish Funding Council [HR03006]. Genotyping of the GS:SFHS samples was carried out by the Genetics Core Laboratory at the Wellcome Trust Clinical Research Facility, Edinburgh, Scotland and was funded by the UK's Medical Research Council and the Wellcome Trust. Ethics approval for the study was given by the NHS Tayside committee on research ethics (reference 05/S1401/89). We are grateful to all the families who took part, the general practitioners and the Scottish School of Primary Care for their help in recruiting them, and the whole Generation Scotland team, which includes interviewers, computer and laboratory technicians, clerical workers, research scientists, volunteers, managers, receptionists, healthcare assistants and nurses. This research has also been conducted using the UK Biobank Resource.

HCW is supported by a JMAS SIM fellowship from the Royal College of Physicians of Edinburgh and by an ESAT College Fellowship from the University of Edinburgh. Part of the work was undertaken in The University of Edinburgh Centre for Cognitive Ageing and Cognitive Epidemiology (CCACE), part of the cross council Lifelong Health and Wellbeing Initiative (MR/K026992/1); funding from the Biotechnology and Biological Sciences Research Council (BBSRC) and Medical Research Council (MRC) is gratefully acknowledged. Age UK (The Disconnected Mind project) also provided support for the work undertaken at CCACE.

## Notes

**Conflicts of Interest:** AMM has previously received grant support from Pfizer, Lilly and Janssen. These studies are not connected to the current investigation. SML has previously received research grant support from Pfizer, Roche, Abbvie and Sunovion, as well as personal fees from Roche, Sunovion and Janssen, outside the present work. Remaining authors report no conflicts of interest.

